# A Persistence Detector for Metabolic Network Rewiring in an Animal

**DOI:** 10.1101/382507

**Authors:** Jote T. Bulcha, Gabrielle E. Giese, Md. Zulfikar Ali, Yong-Uk Lee, Melissa D. Walker, Amy D. Holdorf, L. Safak Yilmaz, Robert C. Brewster, Albertha J.M. Walhout

## Abstract

Biological systems must possess mechanisms that prevent inappropriate responses to spurious environmental signals. Gene regulatory network circuitries known as coherent type 1 feed-forward loops (FFLs) with AND-logic gates have been proposed to function as a persistence detector because it generates a delay in target activation and prevents target induction unless the input signal is sustained. While such a circuit has been found for the L-arabinose utilization system in *E. coli*, their existence and relevance multicellular organisms has remained unclear. Here, we identify the first persistence detector in an animal that redirects propionate breakdown to a shunt pathway when flux through the canonical propionate breakdown pathway is perturbed. We propose that this mechanism has evolved to ensure the shunt pathway stays off unless propionate accumulation is persistent because the shunt pathway generates highly toxic acrylate. Our study uniquely connects persistence detector circuitry to a physiological response in an animal.

## INTRODUCTION

In classic systems biology studies dating more than 15 years ago, Uri Alon and colleagues discovered network motifs, which are defined as small circuits that occur more frequently in different molecular networks than expected by chance (Milo et al., 2002; Shen-Orr et al., 2002). Several network motifs are found in gene regulatory networks that are composed of transcription factors (TFs) and their target genes. One such motif is the feed forward loop (FFL) comprising two TFs each of which regulates a set of downstream targets and one of which modulates the expression or activity of the other (Mangan and Alon, 2003). There are eight types of FFLs, four coherent in which both paths leading to target gene expression are activating or repressing, and four incoherent in which one path is activating and the other is repressing. Modeling these circuitries with different types of end gates such as “AND” where both regulators are required, or “OR” when either one suffices, led to great insights into how these motifs may modulate downstream gene expression. For instance, type 1 coherent FFLs where all regulatory edges are activating coupled with an AND-logic gate, *i.e*., where both TFs are required for downstream target activation, have been proposed to generate a delay in downstream target gene expression because it takes time for the first TF to activate the second (Mangan and Alon, 2003). A consequence of this is that a short input pulse would not be sufficient to activate the second TF in the circuit and therefore fails to induce downstream target gene expression. Consistent with this behavior, coherent type 1 FFLs with an AND-logic gate have been named “persistence detectors” because it takes a sustained, or persistent input for downstream target activation to ensue. While FFLs have been described in many gene regulatory networks, only few persistence detectors have been described, and only in bacteria (Alon, 2007; Lim et al., 2013), and, therefore, their existence and relevance to metazoan biology has remained unclear.

Like humans, the nematode *Caenorhabditis elegans* uses the micronutrient vitamin B12 as a cofactor for two metabolic enzymes: methionine synthase that converts homocysteine into methionine in the methionine (Met)/S-adenosylmethionine (SAM) cycle, and methylmalonyl-CoA mutase, the third enzyme in the canonical propionate breakdown pathway (Watson et al., 2015)(**Figure 1**). Vitamin B12 is exclusively made by bacteria and archaea, and is not used by yeast, plants and flies (Martens et al., 2002). *C. elegans* is a bacterivore, making it a highly suitable “simple” model organism to study vitamin B12 biology (Yilmaz and Walhout, 2014; Zhang et al., 2017). For instance, we have previously found that bacterial diets high in vitamin B12 accelerate *C. elegans* development and change metabolic gene expression (MacNeil et al., 2013; Na et al., 2018; Watson et al., 2014). One set of genes repressed by vitamin B12 comprise an alternative propionate breakdown pathway, or propionate shunt (Watson et al., 2016)(**Figure 1**). Genes in the shunt pathway are activated by excess propionate, which accumulates when flux through the canonical, vitamin B12-dependent propionate breakdown pathway is reduced. This can occur either when the animal eats a bacterial diet low in vitamin B12, or when genes in the canonical pathway are genetically perturbed (Watson et al., 2013; Watson et al., 2014; Watson et al., 2016). The ability to utilize either of two propionate breakdown pathways, depending on metabolic or dietary status, likely provides the animal with metabolic plasticity to thrive on a variety of bacterial diets, be it high or low in vitamin B12. However, the occurrence of two parallel pathways begs the question of why the animal has maintained the canonical propionate breakdown pathway. In other words, why not just use the propionate shunt? The first step in the propionate shunt involves the conversion of propionyl-CoA into acrylyl-CoA, which can be converted into the highly toxic intermediate acrylate upon dissociation of the CoA. Animals in which *ech-6*, encoding the enzyme that metabolizes acrylyl-CoA (**Figure 1**), is perturbed are very sick, whilst a double perturbation of *ech-6* and *acdh-1*, encoding the enzyme that generates acrylyl-CoA rescues this sickness, as the perturbation of *acdh-1* prevents the production of acrylate (Watson et al., 2016). These observations together suggest that the animal has tight control mechanisms to keep the propionate shunt pathway off unless it is needed, such as under persistently low vitamin B12 conditions.

**Figure 1.**
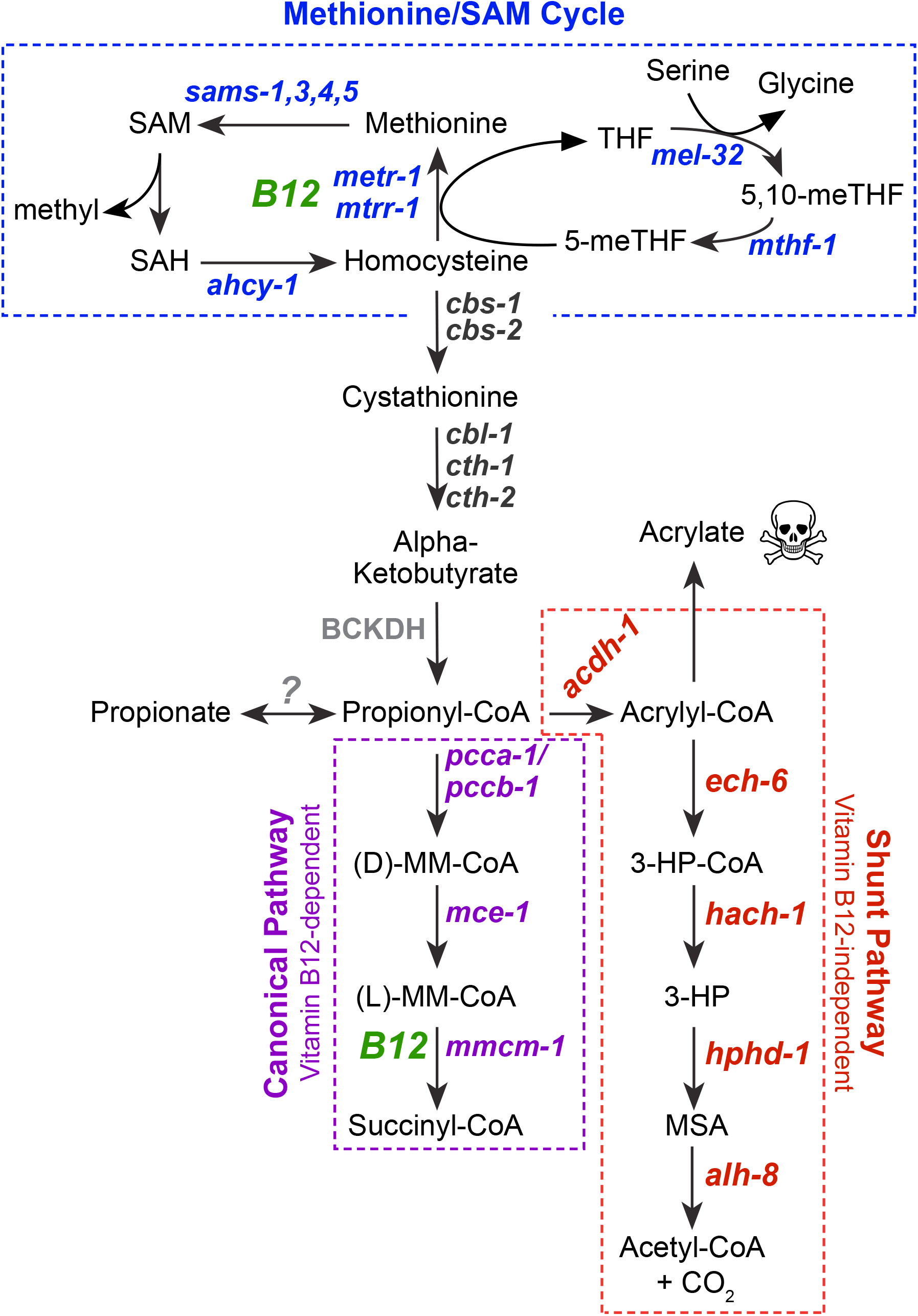
Cartoon of *C. elegans* Metabolic Network Involving Vitamin B12. B12 – vitamin B12, BCKDH - branched-chain ketoacid dehydrogenase, 3-HP – 3-hydroxypropionate, 5,10-meTHF - 5,10-methylenetetrahydrofolate, 5-meTFH – 5-methylenetetrahydrofolate, MM-CoA – methylmalonyl-CoA, MSA – malonic semialdehyde, SAM – S-adenosylmethionine, SAH – S-adenosylhomocysteine, THF – tetrahydrofolate.

Here, we discover a coherent type 1 FFL with an AND-logic gate, composed of two nuclear hormone receptors, in which *nhr-10* activates the expression of *nhr-68*, and both are required for the activation of propionate shunt genes in response to excess propionate. We demonstrate that this circuit functions as a persistence detector in several ways. First, there is a ~3-hour delay in propionate shunt activation upon the supplementation of propionate. Second, a 1-hour pulse of propionate is not sufficient to activate propionate shunt gene expression. Finally, we show that NHR-68 overexpression is not sufficient to activate propionate shunt gene expression in response to propionate, demonstrating that the two NHRs do not function in a simple linear pathway. We propose that the propionate persistence detector functions to ensure that the propionate shunt stays off unless propionate accumulation is persistent, thereby preventing the unwanted generation of highly toxic shunt intermediates. This gene regulatory network architecture links dietary input to metabolic output to ensure animal homeostasis.

## RESULTS

### The Nuclear Hormone Receptors *nhr-10* and *nhr-68* Activate Propionate Shunt Gene Expression in Response to Propionate

We previously generated a *C. elegans* transgenic strain that expresses the green fluorescent protein (GFP) under the control of the *acdh-1* gene promoter (Arda et al., 2010). When these animals are fed an *E. coli* diet that is low in vitamin B12, GFP expression is high, while GFP levels are low on diets high in vitamin B12 (MacNeil et al., 2013; Watson et al., 2014). In addition, GFP expression is activated when genes in the canonical propionate breakdown pathway are genetically perturbed, even in the presence of vitamin B12, or when propionate is supplemented to high-vitamin B12 bacterial diets (Watson et al., 2013). We have previously used the *Pacdh-1::GFP* strain in the context of defining a *C. elegans* intestinal gene regulatory network by comprehensive TF RNAi, and found more than 40 TFs that activate the *acdh-1* promoter on *E. coli* HT115 bacteria (MacNeil et al., 2015). However, it is not clear whether all or only a subset of these TFs specifically mediate the transcriptional response to propionate.

To identify the TF(s) that activate *acdh-1* expression in response to propionate, we tested the TFs previously found on the *E. coli* HT115 diet by RNAi in the presence of both vitamin B12 (to repress the basal GFP expression) and propionate (**Figure S1A**). Interestingly, we found that RNAi of only a subset of the 43 TFs that affected GFP expression on *E. coli* HT115 bacteria also affected GFP levels on propionate supplemented conditions (**Figure 2A, Figure S1B**). Specifically, RNAi of 16 TFs reduced GFP expression under both conditions, RNAi of 26 TFs only repressed GFP expression on untreated conditions, and RNAi of one TF, *mxl-3*, reduced GFP expression on untreated conditions but activated the *acdh-1* promoter on propionate-supplemented conditions. These results indicate that the *acdh-1* promoter not only responds to propionate, but to other cellular conditions as well, and that the response to these other conditions involves other TFs.

**Figure 2.**
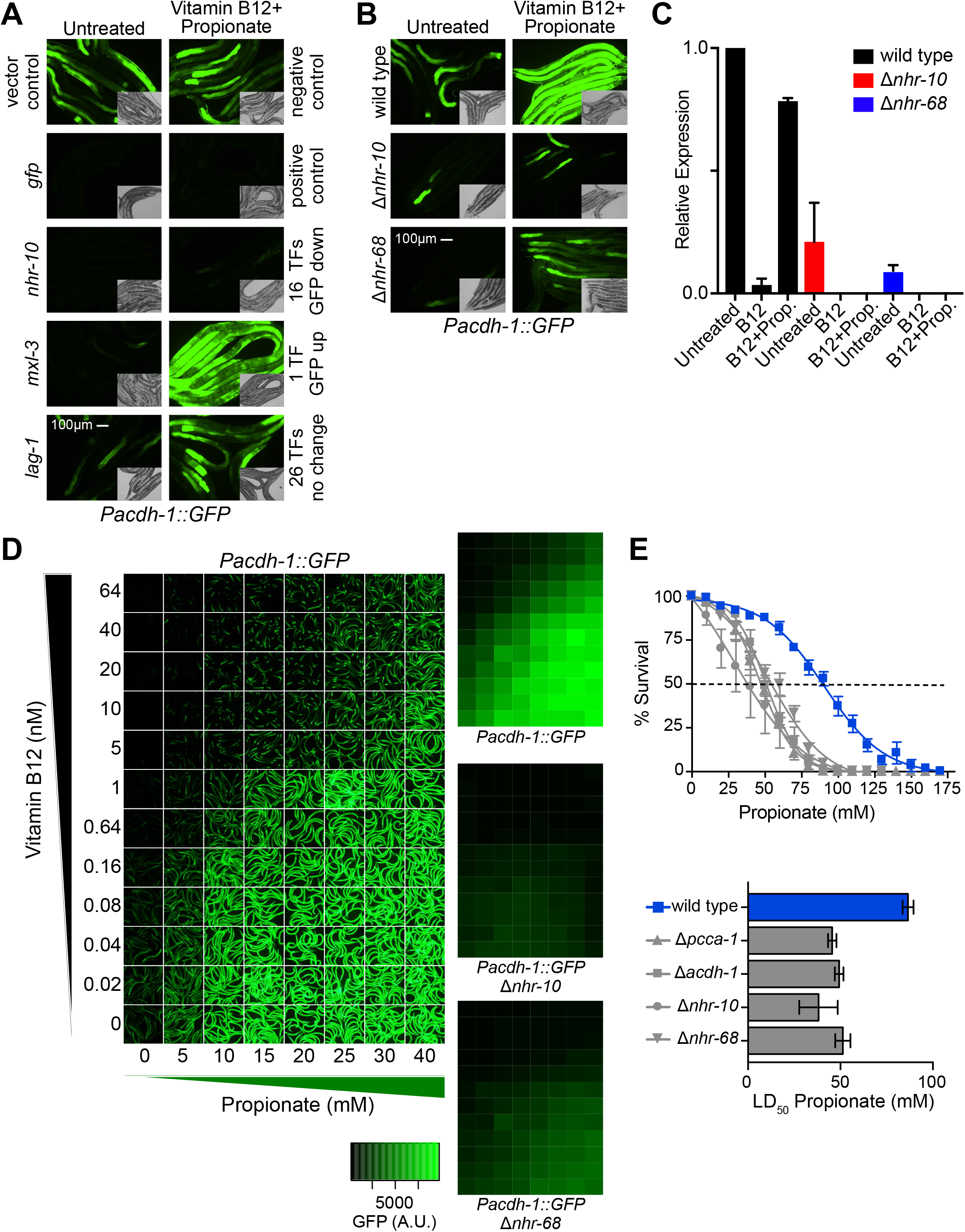
The Nuclear Hormone Receptors NHR-10 and NHR-68 Mediate the Transcriptional and Functional Response to Excess Propionate. (A) Fluorescent microscopy images of TF RNAi shows that only a subset of TFs that activate the *acdh-1* promoter in animals fed *E. coli* HT115 bacteria are involved in the transcriptional response to excess propionate. Insets show DIC images. (B) The transcriptional response to excess propionate is greatly reduced in *nhr-10* and *nhr-68* deletion mutants, validating the TF RNAi results. (C) qRT-PCR experiment showing that endogenous *acdh-1* expression is not induced in response to propionate in *nhr-10* and *nhr-68* deletion mutants. (D) *nhr-10* and *nhr-68* are required for *acdh-1* promoter activation in a broad range of vitamin B12 and propionate concentrations. Left panel shows images of *Pacdh-1::GFP* animals supplemented with indicated concentrations of vitamin B12 and/or propionate. Right three panels show quantification of GFP levels at each concentration in *Pacdh-1::GFP* animals in wild type, *nhr-10* or *nhr-68* deletion mutant animals. Quantification is the average of nine experiments: three biological replicates with three technical replicates each. (E) Propionate toxicity assays show that *nhr-10* and *nhr-68* are functionally required to mitigate the toxic effects of propionate. Top panel shows propionate dose-response curves, bottom panel shows LD_50_ values.

Several of the 16 TFs that reduce GFP expression when knocked down by RNAi function at high levels in the intestinal gene regulatory network and likely affect *acdh-1* promoter activity indirectly (MacNeil et al., 2015). For instance, the intestinal master regulator ELT-2 broadly controls intestinal gene expression and resides at the top of the hierarchy (MacNeil et al., 2015; McGhee et al., 2007). Only two TFs, NHR-10 and NHR-68, have a strong effect on *acdh-1* promoter activity under propionate supplemented conditions and reside low in the gene regulatory network hierarchy (MacNeil et al., 2015). This suggests that they may be critical for propionate shunt activation. Indeed, deletion in either *nhr-10* or *nhr-68* greatly reduced *acdh-1* promoter activity, as well as endogenous *acdh-1* expression (**Figures 2B, 2C**). Quantitative analysis of GFP expression revealed that *nhr-10* and *nhr-68* are both required for *acdh-1* promoter activation under a broad range of propionate concentrations. However, *nhr-10* is absolutely required, while there is still modest activation of GFP expression in *nhr-68* deletion mutant animals (**Figure 2D**).

We previously found that activation of *acdh-1* expression under low vitamin B12 conditions is important to mitigate the effects of excess propionate (Watson et al., 2016). Specifically, the LD_50_ of propionate in *acdh-1* mutant animals fed an *E. coli* diet low in vitamin B12 is similar to that of *pcca-1* deletion mutants, in which flux through the canonical propionate breakdown pathway is perturbed (**Figures 1, 2E**)(Watson et al., 2016). We found that both *nhr-10* and *nhr-68* deletion mutants are more sensitive to excess propionate (**Figure 2E**). Importantly, deletion of either TF renders the animals equally or more sensitive to propionate as deletion of their transcriptional target *acdh-1*, showing that both of these TFs are functionally important to mitigate propionate toxicity. Altogether, these data show that *nhr-10* and *nhr-68* are both required for transcriptional and functional activation of propionate shunt gene expression.

### *nhr-10* and *nhr-68* Activate All Five Propionate Shunt Genes in Response to Excess Propionate

Next, we performed RNA-seq on wild type, *nhr-10* and *nhr-68* mutant animals under untreated, vitamin B12 only and vitamin B12 + propionate supplemented conditions. We identified 23 genes that in wild type animals are repressed by vitamin B12 and activated by propionate, including four of the five propionate shunt genes (**Figure 3A**). The fifth gene, *alh-8*, behaved the same with an adjusted *P*-value <0.01, but was just below our cut-off of a fold-change greater than or equal to 1.5, (**Figure 3B, Table S1**). Activation of thirteen of these 23 genes required both *nhr-10* and *nhr-68*, including the propionate shunt genes (**Figure 3C**). This indicates that a larger gene battery than just the propionate shunt genes is under control of both NHRs.

**Figure 3.**
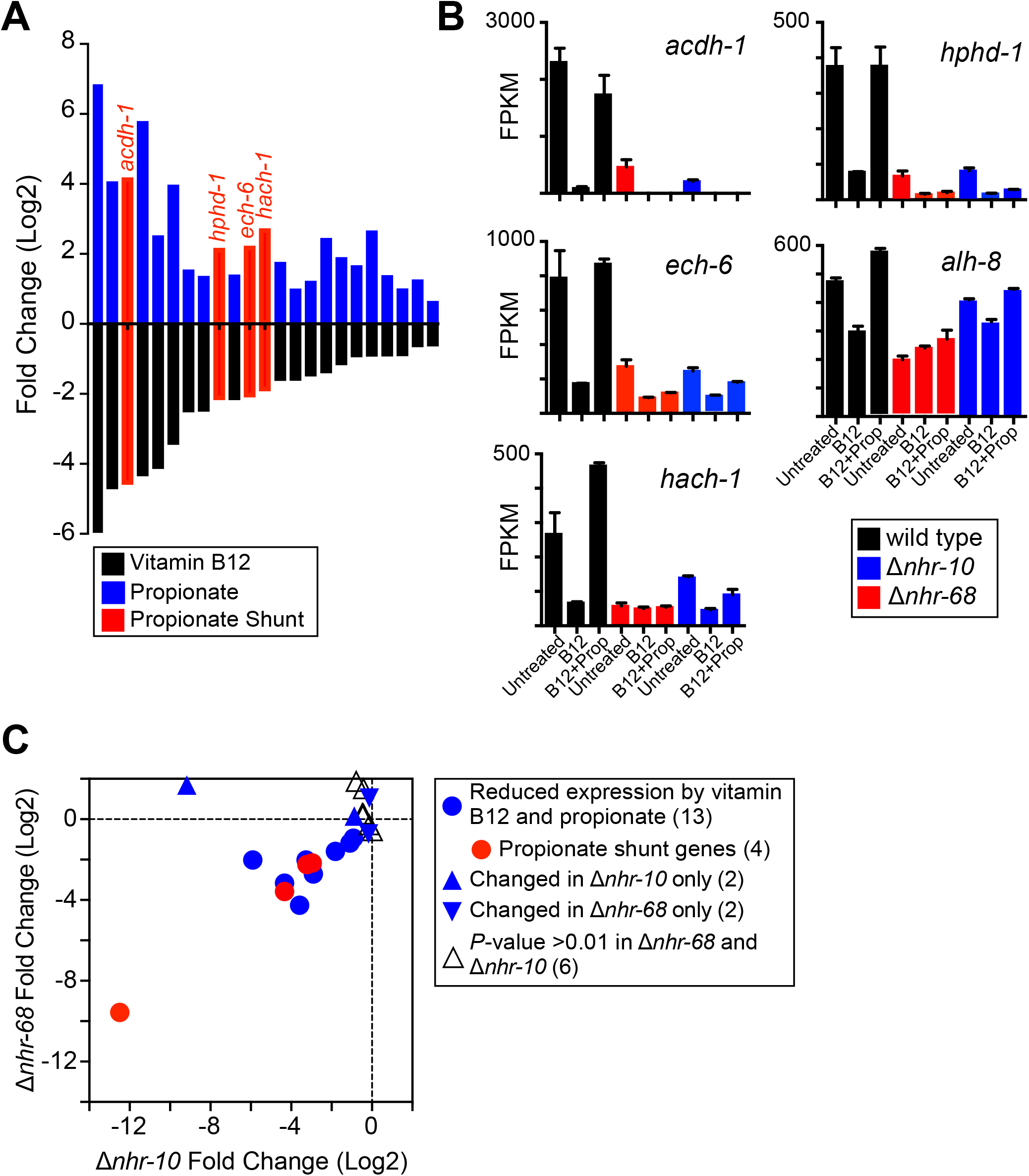
RNA-Seq Expression Profiling of *nhr-10* and *nhr-68* Mutant Animals. (A) Bar-graph showing the 23 genes that are significantly repressed by vitamin B12 and induced by propionate in wild type animals as identified by RNA-seq. (B) Bar-graph of RNA-seq fragments per kilobase of transcript per million mapped reads (FPKM) data showing that all five propionate shunt genes are activated by *nhr-10* and *nhr-68* in response to excess propionate. (C) Scatterplot showing that thirteen of the 23 genes repressed by vitamin B12 and induced by propionate are controlled by both *nhr-10* and *nhr-68*. Gene numbers for each condition in parentheses.

### *nhr-10* and *nhr-68* Function in a Coherent Type 1 FFL with an AND-Logic Gate

While *nhr-10* mRNA levels are not affected by vitamin B12 or propionate supplementation, *nhr-68* is repressed by vitamin B12 and activated by propionate (**Figure 4A, 4B**). Importantly, we found that *nhr-68* mRNA levels are reduced in *nhr-10* deletion mutant animals both by RNA-seq and by qRT-PCR (**Figure 4A, 4B**), which indicates that *nhr-10* activates the expression of *nhr-68*. We validated this result using *Pnhr-68::GFP::H2B* transgenic animals combined with *nhr-10* RNAi which led to a reduction in GFP expression compared to vector control RNAi (**Figure 4C**). Interestingly, *nhr-68* RNAi also caused a reduction in GFP levels in this strain, indicating that *nhr-68* is an auto-activator. Altogether, our observations suggest that *nhr-10* and *nhr-68* function in a coherent type 1 FFL with an AND-logic gate (**Figure 4D**). However, our data so far do not exclude the possibility that *nhr-10* and *nhr-68* function in a simple linear pathway where *nhr-10* activates *nhr-68* and *nhr-68* activates propionate shunt gene expression. To test this idea, we over-expressed *nhr-68* under the control of the promoter of moderately and highly constitutively expressed intestinal genes, *ges-1* and *asp-5*, respectively. Neither of these genes is affected by vitamin B12 or propionate, or by deletion of *nhr-10* (**Figure S2A, S2B**). We examined induction of *acdh-1* expression by qRT-PCR either with vector control or with *nhr-10* RNAi in both strains. We found that, in these animals, *acdh-1* mRNA levels were repressed by vitamin B12 and induced by propionate, just as in wild type animals (**Figure 4E, Figure S2C**). However, *acdh-1* levels were greatly reduced upon RNAi of *nhr-10*, indicating that *nhr-10* is absolutely required for propionate shunt activation. This finding demonstrates that activation of *nhr-68* by *nhr-10* is not sufficient for propionate shunt gene induction in response to propionate, but rather that the path leading from *nhr-10* to these genes is essential as well.

**Figure 4.**
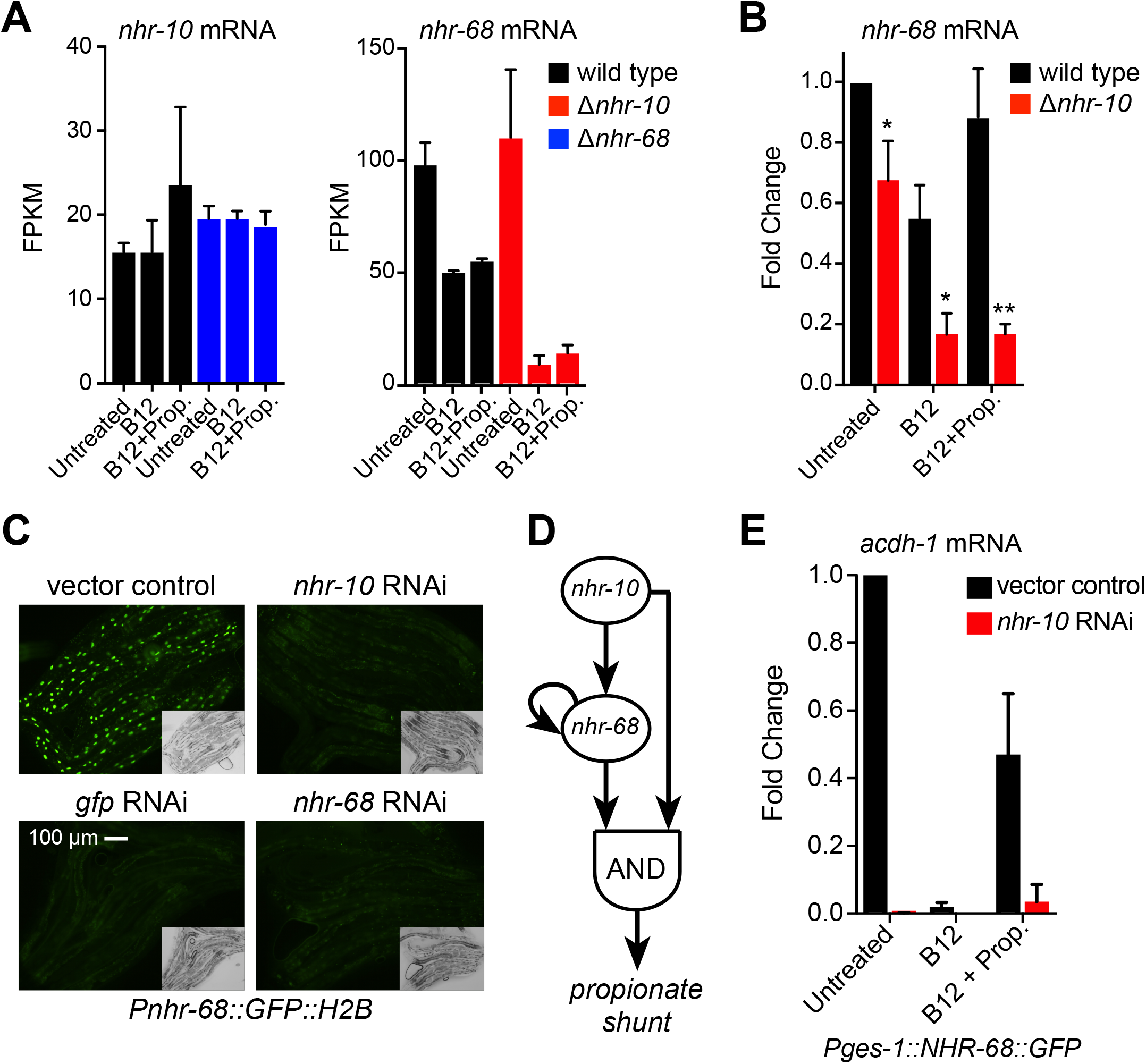
*nhr-10* Activates *nhr-68* Expression and *nhr-68* is an Auto-Activator. (A) Bar-graph of RNA-seq data for *nhr-10* and *nhr-68* mRNA levels in the absence of *nhr-68* (left) and *nhr-10* (right). (B) Bar-graph showing qRT-PCR data for *nhr-68* mRNA levels in the absence of *nhr-10*. Statistical differences between wild type and *nhr-10* deletion were determined by twotailed paired Student’s t-test (* *P* < 0.05, ** *P* < 0.005). (C) Fluorescent microscopy images of RNAi of *nhr-10* or *nhr-68* shows reduced GFP expression in *Pnhr-68::GFP::H2B* transgenic animals. Insets show DIC images. (D) Cartoon of the propionate persistence detector. (E) qRT-PCR shows that constitutive intestinal expression (*ges-1* promoter) of NHR-68 does not induce *acdh-1* expression in response to propionate. All measurements are statistically significantly different compared to untreated vector control as determined by two-tailed paired Student’s t-test (P<0.05).

### The *nhr-10/nhr-68* Coherent Type 1 FFL with an AND-Logic Gate Functions as a Propionate Persistence Detector

The finding that *nhr-10* and *nhr-68* function in a coherent type 1 FFL with an AND-logic gate leads to the prediction that this circuit may function as a propionate persistence detector that ensures that propionate shunt gene expression is only induced when propionate is present sufficiently long, and that this gene activation occurs with a delay (Alon, 2007). To test this prediction, we first constitutively supplemented *Pacdh-1::GFP* transgenic animals kept on vitamin B12 (GFP expression off) with propionate and quantified GFP expression every 30 minutes the first two hours and every hour for 22 hours, and after 30 and 36 hours. As expected, GFP stayed off in animals that were constitutively supplemented with vitamin B12 (**Figure 5A**). When supplemented with propionate, however, GFP expression was robustly induced. Importantly, this induction occurred only after ~3 hours of propionate supplementation, confirming a delay in propionate shunt gene expression that would be expected for a true persistence detector. Second, we found that when *Pacdh-1::GFP* animals were given the same concentration of propionate, but only in a 1-hour pulse, GFP expression was not induced (**Figure 5B**). These findings further support the model that *nhr-10* and *nhr-68* act in a coherent type 1 FFL with an AND-logic gate that serves to function as a persistence detector in an animal.

**Figure 5.**
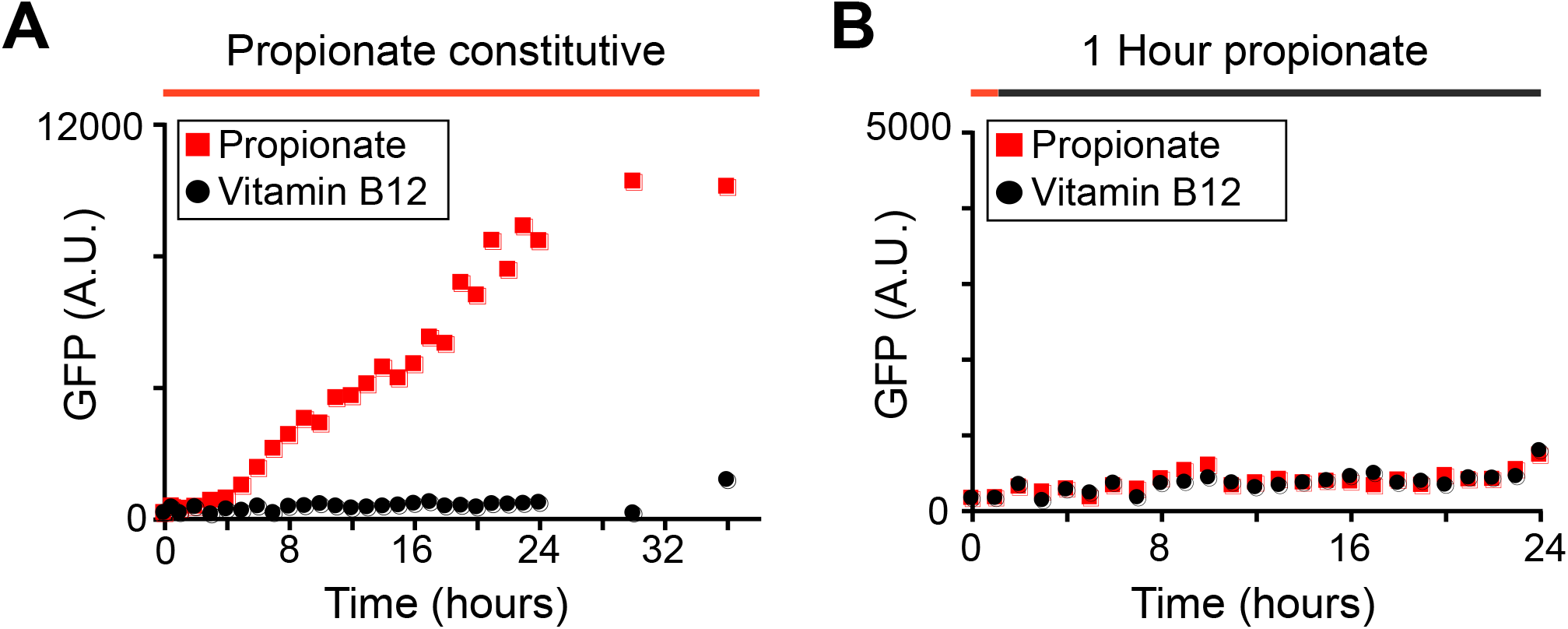
*acdh-1* Expression is Induced with a Three-Hour Delay in Response to Propionate and does not Respond to a One-Hour Propionate Pulse. (A) GFP expression in *Pacdh-1::GFP* animals transferred from vitamin B12 (black circles) to constitutive propionate supplementation (red squares) shows a ~three hour delay. (B) A one-hour pulse of propionate does not induce GFP expression in *Pacdh-1::GFP* animals.

### *nhr-68* Autoactivation can Modulate the Delay in Persistence Detector Target Gene Activation

Next, we asked how *nhr-68* auto-activation may contribute to the functionality of the persistence detector. For this, we modeled a type 1 coherent FFL with an AND-gate with autoregulation of *nhr-68* and simulated the system using Michaelis-Menten kinetics. We used the fitted experimental data from **Figure 5A** to estimate the parameters (**Figure 6A**, see Experimental Procedures). We then explored the interplay between *nhr-68* basal expression rate (characterized by the parameter, *r)* and *nhr-68* autoregulatory expression rate (characterized by *r_A_)* on the observed delay in target gene expression. **Figure 6B** shows a heat map of delay times as a function of these two parameters over a broad range of values. Although precise quantitative values of the model parameters are not known, we found an interesting general feature of the circuit. Without auto-activation, the system has a tight range of predicted delay times that is relatively insensitive to the basal rate of induction and is set by the decay rates of the constituent TFs (**Figure 6C**). However, the inclusion of autoregulation of *nhr-68* enables a wide range of delays that are set by the basal expression rate (which, in our model, is controlled solely by *nhr-10*). In other words, with auto-activation, small basal rates resulting from lower input signals would trigger longer delays in target gene expression while strong input signals would trigger shorter delays and a quicker response (**Figure 6C, 6D**).

**Figure 6.**
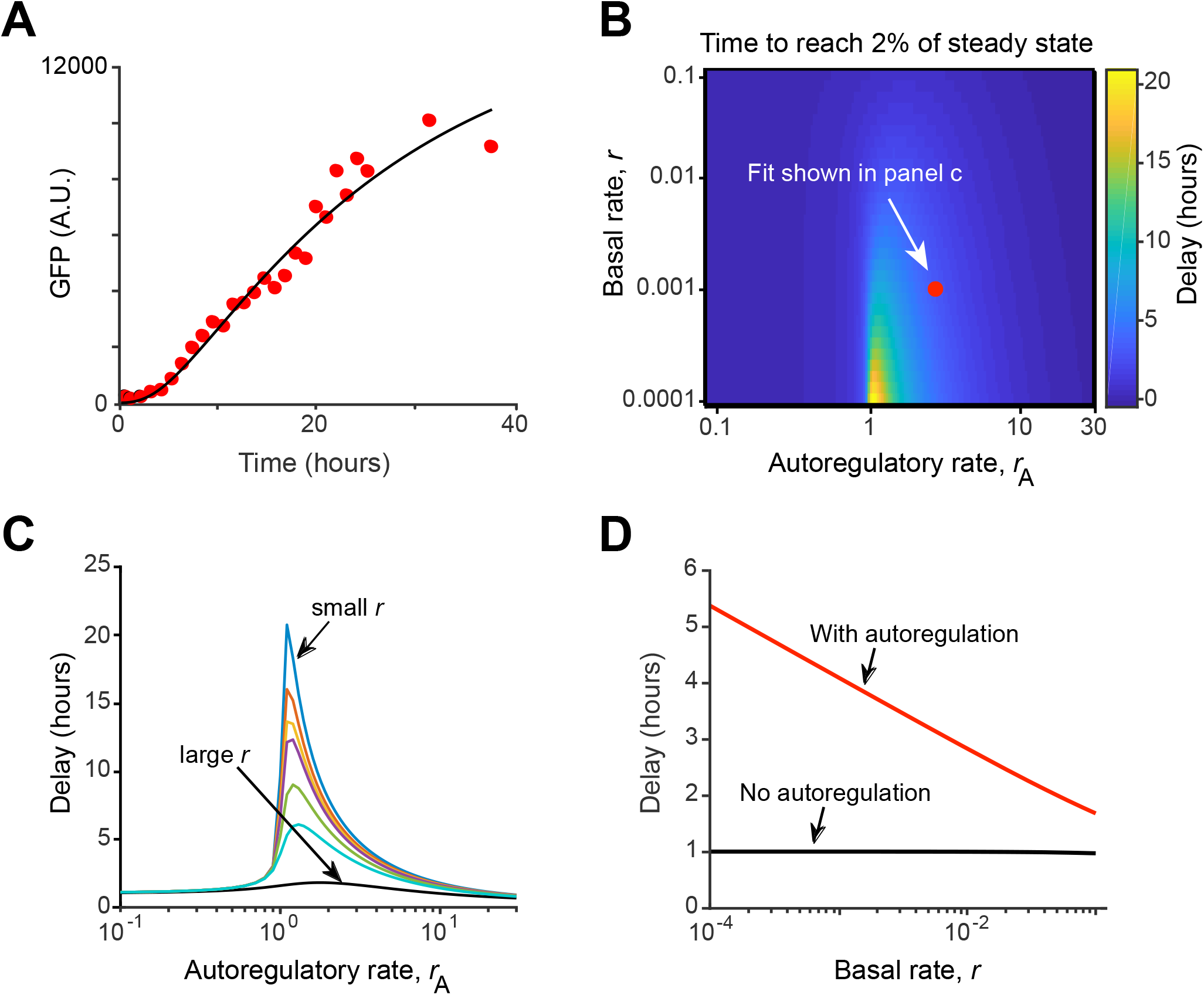
Computational Modeling Shows that *nhr-68* Autoactivation Can Modulate the Delay in Target Gene Expression. (A) Fit of propionate supplementation data from Figure 5A to estimate approximate parameter range. (B) Heat map of delay time to reach 2% of steady state level of GFP based on the rate of basal expression, *r*, and the rate of autoregulation, *r_A_*, of the *nhr-68* gene. (C) Delay time as a function of autoregulatory strength for several basal rates from the range of values examined in (B). (D) Delay time with autoregulation (using *r_A_* found from C) and without autoregulation as basal rate is tuned.

## DISCUSSION

We have discovered the first persistence detector in an animal. This persistence detector activates the five genes comprising the propionate shunt pathway in *C. elegans*. This pathway provides an alternative way to catabolize propionate, which is toxic when it accumulates in both humans and *C. elegans* (Deodato et al., 2006; Watson et al., 2016). Shunting of propionate occurs in both humans and *C. elegans*, as indicated by the detection of shunt pathway intermediates in propionic acidemia patients and individuals with mutations in other relevant genes (Ando et al., 1972; Deodato et al., 2006; Peters et al., 2015). However, while *C. elegans* has evolved a propionate shunt pathway that is transcriptionally induced when propionate accumulates, humans likely repurpose metabolic enzymes that function in other metabolic pathways (Watson et al., 2016).

Two observations indicate that *C. elegans* preferentially utilizes the canonical, vitamin B12-dependent propionate breakdown pathway, rather than the propionate shunt. First, the canonical pathway has not been lost in evolution, which would be expected if its function became obsolete. Second, the shunt pathway is inactive when flux through the canonical pathway is enabled by the sufficient dietary intake of vitamin B12. One reason for preferential use of the canonical pathway is that the propionate shunt generates acrylate, which is more toxic than propionate. Acrylate is produced in the first reaction in the propionate shunt where propionyl-CoA is converted into acrylyl-CoA (which can be interconverted with acrylate when the CoA can be chemically or enzymatically removed) by the ACDH-1 enzyme (**Figure 1**). The next enzyme in the shunt pathway, ECH-6, then converts acrylyl-CoA into 3-hydroxypropionyl-CoA. We previously found that RNAi of *ech-6* renders the animals very sick, and that double perturbation of *ech-6* with *acdh-1* suppresses this phenotype (Watson et al., 2016). Our data indicate that *C. elegans* ensures that the shunt pathway stays off until it is really needed by utilizing a transcriptional persistence detection mechanism.

The propionate persistence detector consists of two NHRs: *nhr-10* and *nhr-68*, where *nhr-10* activates *nhr-68, nhr-68* auto-activates and both activate shunt gene expression in response to persistent propionate accumulation. While our study illuminates the systems-level mechanism of propionate persistence detection, the precise molecular mechanism remains to be elucidated. We previously found that NHR-10 physically binds the *acdh-1* promoter in yeast one-hybrid assays (Arda et al., 2010; MacNeil et al., 2015). We do not yet know whether it also interacts with the other shunt gene promoters. One possibility is that NHR-68 and NHR-10 physically interact to form a heterodimer. However, we did not detect any interactions with NHR-68 in our large-scale protein-DNA and protein-protein interaction studies (Fuxman Bass et al., 2016; Reece-Hoyes et al., 2013), so this remains to be investigated. We also do not yet know the mechanism of propionate detection by the persistence detector. Propionate is generated from odd-chain fatty acids, branched-chain amino acids, methionine and threonine (Yilmaz and Walhout, 2016), and is catabolized in the mitochondria (Al-Lahham et al., 2010). How information involved in propionate shunt activation is transferred from the mitochondria and cytoplasm to NHRs in the nucleus is not known. It is tempting to speculate that NHR-10, NHR-68, or both are directly activated by propionate, which is a three-carbon short chain fatty acid, given that NHRs are known to use fatty acids as ligands (Evans and Mangelsdorf, 2014). However, the small size and volatility of propionate make detection of its interaction with proteins extremely challenging.

Several observations indicate that the persistence detector does not function in isolation and does not function solely to activate the expression of *acdh-1* and other genes, but rather, that it is embedded in a larger gene regulatory network. First, the set of downstream targets consists of at least 13 genes that are repressed by vitamin B12, activated by propionate and dependent on both *nhr-10* and *nhr-68*. Aside from four shunt genes, this set contains nine genes the function of which is largely unknown. It is likely that several of these genes function to support shunt function or to enable its shutdown when nutritional conditions favor the use of the canonical vitamin B12-dependent propionate breakdown pathway. Second, many additional TFs are involved in *acdh-1* expression ((MacNeil et al., 2015), this study). These include 16 TFs that, when knocked down by RNAi, reduce *acdh-1* promoter activity either induced by propionate supplementation or under untreated conditions on the *E. coli* HT115 diet. Most of these TFs have a partial effect and reside higher in the intestinal gene regulatory network (MacNeil et al., 2015) and likely affect the *acdh-1* promoter indirectly. RNAi of one TF, *mxl-3*, reduced *acdh-1* promoter activity on *E. coli* HT115 bacteria but increased it under propionate supplemented conditions. We did not follow up on this observation because we observed only very small effects on propionate shunt gene expression in *mxl-3* mutant animals (data not shown). Another set of 26 TFs regulate *acdh-1* under untreated conditions only, *i.e*., their knockdown has no effect on propionate supplemented conditions. This finding indicates that *acdh-1* responds to other metabolites that act via other TFs. Interestingly, these TFs include *nhr-101* and *nhr-114*, which would be appealing candidates to mediate the response to other metabolites. It is interesting to note that *acdh-1* expression is not completely off in either *nhr-10* or *nhr-68* deletion mutant in the untreated condition on *E. coli* HT115 (**Figure 1B**) and that this residual expression is repressed by vitamin B12. This observation suggests that other metabolites activating *acdh-1* may also be functionally connected to vitamin B12 metabolism. Moreover, it suggests that *acdh-1* may have a function outside of the propionate shunt. Taken together, we discovered the first persistence detector and link this gene regulatory network architecture to a functional metabolic response in a whole animal.

## MATERIALS AND METHODS

### *C. elegans* and *E. coli* Strains

N2 (Bristol) was used as the wild type strain, and animals were maintained on nematode growth medium (NGM) at 20°C as described (Brenner, 1974). Strains VL1286 (wwSi28[*Pnhr-68::GFP::H2B*; *unc-119*(+) II]), VL1296 (wwSi29[*Pges-1::NHR-68::GFP*; *unc-119*(+) II]), VL1297 (wwSi30[*Pasp-5::NHR-68*; *unc-119*(+) II]) were constructed by Gateway cloning (ThermoFisher Scientific) using DNA from the *C. elegans* promoterone (Dupuy et al., 2007), and transgenic strains were made using the *Mos1*-mediated single-copy insertion (MosSCI) method (Frokjaer-Jensen et al., 2014). Integrated transgenes were confirmed by PCR genotyping (primers for genotyping in **Table S2**). *C. elegans* strains *nhr-68(gk708)*, *pcca-1(ok2282)*, *acdh-1(ok1489)* and *E. coli* strains OP50 and HT115 were provided by the *Caenorhabditis* Genetics Center (CGC). *C. elegans* strains *nhr-10(tm4695)* was provided by the National Bioresource Project, Japan. *C. elegans* strains VL749 (wwIs24[*Pacdh-1::GFP*; *unc-119*(+)]) and VL868 (*nhr-10(tm4695)*; wwIs24[*Pacdh-1::GFP*; *unc-119*(+)]) were previously described (MacNeil et al., 2013), and *C. elegans* strain VL1113 (*nhr-68(tm708)*; wwIs24[*Pacdh-1::GFP*; *unc-119*(+)]) was previously described (Watson et al., 2013). *E. coli* HT115 RNAi strains were obtained from the Ahringer (Kamath et al., 2003) or ORFeome (Rual et al., 2004) RNAi libraries.

### RNA Interference

RNAi was performed as described (MacNeil et al., 2015) with or without supplementation of 5 nM vitamin B12 and 40 mM propionate (propionic acid, Sigma Aldrich). Changes in intestinal GFP were scored visually when samples contained a mix of L4 and young adult animals. Knockdowns were scored as positive when most animals in the well displayed a change in intestinal GFP. Changes in GFP levels in other tissues were not recorded. Experiments were performed five independent times. TFs that scored in at least three independent experiments were considered hits. All RNAi clones included in the final dataset were sequence-verified.

### Propionate and Vitamin B12 Gradient Assays

L1 synchronized animals were grown on NGM media supplemented with a matrix of propionate (propionic acid, Sigma Aldrich, 0 to 40 mM) and vitamin B12 (adenosyl cobalamin, Sigma Aldrich, 0 to 64 nM) concentrations. GFP fluorescence was measured using a Tecan Infinite M1000Pro microplate reader as described (Leung et al., 2011). Five adult animals were randomly picked in triplicate and transferred to a 384 well plate containing 35 µl of M9 buffer containing 1 µM levamisole (levamisole hydrochloride, Sigma Alrich) and 0.5% Polyethylene Glycol (PEG, Sigma Aldrich). GFP intensity measurement was performed at 485nm/9nm excitation and 535nm/20nm emission spectra. Each experiment was performed in biological triplicate with three technical triplicates each, and the fluorescence intensity of each biological replicate was averaged. The average fluorescence intensity was used to make the heatmap using the online tool http://www.heatmapper.ca/expression/ (Babicki et al., 2016).

### Propionate Toxicity Assays

Propionate toxicity assays were performed as described previously (Watson et al., 2016). Approximately 100-200 synchronized L1 animals (hatched overnight, 20 hours post-bleach) were added to *E. coli* OP50-seeded 35 mm NGM agar plates containing various concentrations of pH-neutralized propionic acid. Each dose tested included two technical and three biological replicates. After 72 hours, animals that had developed past L1 stage were counted. The fraction surviving, *S*, as a function of propionic acid concentration, *[C]*, was fit to the following dose response curve:

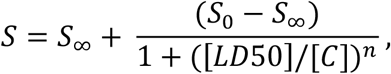

where *S_0_* and *S_∞_* are the fraction of animals surviving at zero and at saturating concentrations of propionic acid and *n* is the Hill slope. The dose required to kill 50% of the population, LD_50_, was found by least-squares fit of these four parameters. Toxicity assays were performed in biological triplicate, and the average LD_50_ was plotted ± SEM.

### Expression Profiling

Animals were fed *E. coli* OP50 on NGM-agar supplemented with 10 nM vitamin B12. Animals were then grown on NGM-agar without vitamin B12 and synchronized for two generations by L1 arrest in M9 buffer for 18 hours post-treatment with buffered bleach. Animals were then grown on NGM-agar alone, or on NGM-agar supplemented with either 20 nM vitamin B12 or 20 nM vitamin B12 and 40 mM propionate. Approximately 3000 L4 stage animals were harvested and washed three times in M9 buffer for each condition. Total RNA was isolated using Trizol (Life Technologies) followed by DNase I (NEB) treatment and cleanup using Direct-zol RNA Mini Prep kit (Zymo Research). Two biological replicates for each condition were sequenced. Sequencing was performed by BGI using BGISEQ-500 platform with a single-end 50 bp read length, and a minimum of 26 million reads per sample. Differential gene expression between samples was analyzed using the standard output of BGI bioinformatics pipeline using EBSeq (Leng et al., 2013) for identifying differentially expressed genes. Differentially expressed genes were selected based on a fold change of ≥ ± 1.5, and adjusted *P*-value ≤ 0.01.

### qRT-PCR

qRT-PCR was performed as described previously (Watson et al., 2016). Briefly, synchronized L1 animals were grown on NGM-agar plates containing 20 nM vitamin B12 and/or 40 mM propionate seeded with *E. coli* OP50 and grown at 20^°^C until they reached to late L4 stage. About 1500 animals were harvested for each condition, in triplicate. Aniamls were washed in M9 buffer and total RNA was isolated using TRIzol Reagent (Life Technologies), following by DNAseI (NEB) treatment and cleanup with Direct-zol RNA Mini Prep Kit (Zymo Research). cDNA was prepared from RNA using Oligo(dT) 12-18 Primer (Invitrogen), RNAseOut (Invitrogen), and M-MuLV Reverse Transcriptase (NEB). qPCR was performed in technical triplicate per gene condition using the Applies Biosystems StepOnePlus Real-Time PCR system and Fast Sybr Green Master Mix (ThermoFisher Scientific). Relative transcript abundance was determined by using the ΔΔCt method (18546601) and normalized to averaged *ama-1* and *act-1* mRNA expression levels. Primer sequences are provided in **Table S2**.

### Propionate Pulse Experiments

L1 synchronized animals were grown on 10 nM vitamin B12 supplemented media seeded with *E. coli* OP50 until the adult stage. Animals were then transferred to 40 mM propionate supplemented media either constitutively, or for one hour, and transferred back to vitamin B12 supplemented media. During each transfer the animals were washed three times using M9 buffer. For each time point, five adult animals were randomly picked and transferred to a 384 well plate containing 35 µl M9 buffer containing 1 µM levamisole and 0.5% PEG. GFP was measured at 485nm/20nm excitation and 535nm/20nm emission spectra as described above. Each experiment was performed in biological triplicate with three technical replicates each, and the fluorescence intensity of technical replicates were averaged for each biological replicate.

### Modeling FFL with Positive Autoregulation

In the coherent FFL motif with an AND-logic gate (**Figure 4D**), propionate causes *nhr-10* to activate *nhr-68* expression with a basal rate *r*, and both *nhr-10* and *nhr-68* jointly activate GFP expression through an AND logic-gate with a rate *r_c_*. In the absence of the signal, *r_c_* =0 due to the AND logic. Additionally, *nhr-68* auto-activates with rate *r_A_*. Implicit in this model is the assumption that *nhr-10* acts as a switch; *i.e. nhr-10* is “on” in the presence of propionate and “off” in its absence, and that levels of *nhr-10* do not change with time. To model the dynamics of the system we used Michaelis-Menten kinetics to arrive at the following ODEs that describe the system. GFP expression driven by the *acdh-1* promoter in *Pacdh-1::GFP* transgenic animals was used as a proxy for propionate shunt expression:

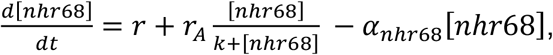

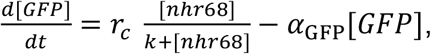

where *α* is the degradation rate of *nhr-68* or GFP and *k* is the dissociation constant. We assumed the degradation rate of *nhr-68* to be relatively fast and set *α_nhr68_* = 1/hour. The decay rate of GFP, which is known to be very stable, was set to *α_GFP_* = 0.05/hour which is determined from fitting the decay of GFP signal over time when propionate is washed out; although *α_GFP_* does not have biological relevance, it impacts the dynamics of our measured signal and the rate of approach to steady state. The dissociation constant is set to 1. The steady state concentration of GFP is obtained by setting the ODEs to zero and solving for GFP and *nhr-68* levels. The delay time, defined as the time to reach 2% of the maximum GFP level, is numerically obtained by solving the ODEs using the ODE solver, ODE45, in MATLAB. We fit the data for *r*, *r_A_* and *r_c_* in order to estimate the magnitude of these parameters. The role of autoregulation in the circuit was tested by varying the two contributions to *nhr-68* production (autoregulation, characterized by *r_A_* and basal expression from *nhr-10* alone, characterized by *r)* around these best-fit values. Adjusting the parameters *k* and *α_nhr68_* alter the fit parameters, however, the highlighted qualitative features of the circuit remain unchanged.

## ACKNOWLEDGEMENTS

We thank members of the Walhout lab and the faculty in the Program in Systems Biology for discussion and critical reading of the manuscript. This work was supported by a grant from the National Institutes of Health DK068429 to A.J.M.W. Some bacterial and nematode strains used in this work were provided by the CGC, which is funded by the NIH Office or Research Infrastructure Programs (P40 OD010440).

## AUTHOR CONTRIBUTIONS

J.T.B. and G.E.G. performed all experiments with technical help from M.D.W. L.S.Y. analyzed the RNA-seq data. Y-U.L. generated the *Pnhr-68::GFP::H2B* construct. A.D.H. helped with analysis, writing and making figures. M.Z.A. did the modeling shown in Figure 6 under supervision of R.C.B. A.J.M.W. conceived the project and wrote the manuscript, with help from all other authors.

---

**Figure S1. Results of the TF RNAi screen for the increase or decrease of GFP expression in *Pacdh-1::GFP* transgenic animals in the presence of propionate. Related to Figure 2.**

(A) Fluorescence microscopy images (bottom) show that 5 nM vitamin B12 represses *Pacdh-1::GFP* while propionate induces reporter activation in a dose dependent manner. Top images show DIC controls.

(B) Fluorescence microscopy images of all 17 TF RNAi experiments performed side-by-side with untreated animals and with animals supplemented with 5 nM vitamin B12 and 40 mM propionate. Insets show DIC images.

**Figure S2: Expression levels of *ges-1* and *asp-5* are not affected by vitamin B12, propionate, *nhr-10*, or *nhr-68*. Related to Figure 4**.

(A, B) Bar-graph of RNA-seq fragments per kilobase of transcript per million mapped reads (FPKM) data showing that expression levels of intestinal genes *ges-1* (low expression) and *asp-5* (high expression) are not affected by 20 nM vitamin B12, 40 mM propionate, *nhr-10* deletion, or *nhr-68* deletion.

(C) qRT-PCR showing that constitutive intestinal expression (*asp-5* promoter) of NHR-68 does not induce *acdh-1* expression in response to propionate. All measurements are statistically significantly different compared to untreated vector control as determined by two-tailed paired Student’s t-test (P<0.05).

**Table S1, related to Figure 3**: RNA-seq fold change and P-adjusted values for the 23 genes up regulated by propionate and down regulated by vitamin B12 in wild type animals.

**Table S2, related to the Experimental Procedures**: All DNA oligonucleotides used in this study.

## REFERENCES

Al-Lahham, S.H., Peppelenbosch, M.P., Roelofsen, H., Vonk, R.J., and Venema, K. (2010). Biological effects of propionic acid in humans; metabolism, potential applications and underlying mechanisms. Biochimica et biophysica acta 1801, 1175-1183.

Alon, U. (2007). Network motifs: theory and experimental approaches. Nat Rev Genet 8, 450-461.

Ando, T., Rasmussen, K., Nyhan, W.L., and Hull, D. (1972). 3-hydroxypropionate: significance of -oxidation of propionate in patients with propionic acidemia and methylmalonic acidemia. Proc Natl Acad Sci U S A 69, 2807-2811.

Arda, H.E., Taubert, S., Conine, C., Tsuda, B., Van Gilst, M.R., Sequerra, R., Doucette-Stam, L., Yamamoto, K.R., and Walhout, A.J.M. (2010). Functional modularity of nuclear hormone receptors in a *C. elegans* gene regulatory network. Molecular Systems Biology 6, 367.

Babicki, S., Arndt, D., Marcu, A., Liang, Y., Grant, J.R., Maciejewski, A., and Wishart, D.S. (2016). Heatmapper: web-enabled heat mapping for all. Nucleic Acids Res 44, W147-153.

Brenner, S. (1974). The genetics of *Caenorhabditis elegans*. Genetics 77, 71-94.

Deodato, F., Boenzi, S., Santorelli, F.M., and Dionisi-Vici, C. (2006). Methylmalonic and propionic aciduria. Am J Med Genet C Semin Med Genet 142C, 104-112.

Dupuy, D., Bertin, N., Hidalgo, C.A., Venkatesan, K., Tu, D., Lee, D., Rosenberg, J., Svrzikapa, N., Blanc, A., Carnec, A., et al. (2007). Genome-scale analysis of in vivo spatiotemporal promoter activity in Caenorhabditis elegans. Nat Biotechnol 25, 663-668.

Evans, R.M., and Mangelsdorf, D.J. (2014). Nuclear Receptors, RXR, and the Big Bang. Cell 157, 255-266.

Frokjaer-Jensen, C., Davis, M.W., Sarov, M., Taylor, J., Flibotte, S., LaBella, M., Pozniakovsky, A., Moerman, D.G., and Jorgensen, E.M. (2014). Random and targeted transgene insertion in *Caenorhabditis elegans* using a modified Mos1 transposon. Nat Methods 11, 529-534.

Fuxman Bass, J.I., Pons, C., Kozlowski, L., Reece-Hoyes, J.S., Shrestha, S., Holdorf, A.D., Mori, A., Myers, C.L., and Walhout, A.J.M. (2016). A gene-centered *C. elegans* protein-DNA interaction network provides a framework for functional predictions. Mol Syst Biol 12, 884.

Kamath, R.S., Fraser, A.G., Dong, Y., Poulin, G., Durbin, R., Gotta, M., Kanapin, A., Le Bot, N., Moreno, S., Sohrmann, M., et al. (2003). Systematic functional analysis of the *Caenorhabditis elegans* genome using RNAi. Nature 421, 231-237.

Leng, N., Dawson, J.A., Thomson, J.A., Ruotti, V., Rissman, A.I., Smits, B.M., Haag, J.D., Gould, M.N., Stewart, R.M., and Kendziorski, C. (2013). EBSeq: an empirical Bayes hierarchical model for inference in RNA-seq experiments. Bioinformatics 29, 1035-1043.

Leung, C.K., Deonarine, A., Strange, K., and Choe, K.P. (2011). High-throughput screening and biosensing with fluorescent *C. elegans* strains. J Vis Exp.

Lim, W.A., Lee, C.M., and Tang, C. (2013). Design principles of regulatory networks: searching for the molecular algorithms of the cell. Mol Cell 49, 202-212.

MacNeil, L.T., Pons, C., Arda, H.E., Giese, G.E., Myers, C.L., and Walhout, A.J.M. (2015). Transcription factor activity mapping of a tissue-specific gene regulatory network. Cell Syst 1, 152-162.

MacNeil, L.T., Watson, E., Arda, H.E., Zhu, L.J., and Walhout, A.J.M. (2013). Diet-induced developmental acceleration independent of TOR and insulin in *C. elegans*. Cell 153, 240-252.

Mangan, S., and Alon, U. (2003). Structure and function of the feed-forward loop network motif. Proc Natl Acad Sci U S A 100, 11980-11985.

Martens, J.H., Barg, H., Warren, M.J., and Jahn, D. (2002). Microbial production of vitamin B12. Appl Microbiol Biotechnol 58, 275-285.

McGhee, J.D., Sleumer, M.C., Bilenky, M., Wong, K., McKay, S.J., Goszczynski, B., Tian, H., Krich, N.D., Khattra, J., Holt, R.A., et al. (2007). The ELT-2 GATA-factor and the global regulation of transcription in the *C. elegans* intestine. Developmental biology 302, 627-645.

Milo, R., Shen-Orr, S., Itzkovitz, S., Kashtan, N., Chklovskii, D., and Alon, U. (2002). Network motifs: simple building blocks of complex networks. Science 298, 824-827.

Na, H., Ponomarova, O., Giese, G.E., and Walhout, A.J.M. (2018). *C. elegans* MRP-5 exports vitamin B12 from mother to offspring to support embryonic development. Cell Rep 22, 3126-3133.

Peters, H., Ferdinandusse, S., Ruiter, J.P., Wanders, R.J., Boneh, A., and Pitt, J. (2015). Metabolite studies in HIBCH and ECHS1 defects: Implications for screening. Mol Genet Metab 115, 168-173.

Reece-Hoyes, J.S., Pons, C., Diallo, A., Mori, A., Shrestha, S., Kadreppa, S., Nelson, J., DiPrima, S., Dricot, A., Lajoie, B.R., et al. (2013). Extensive rewiring and complex evolutionary dynamics in a *C. elegans* multiparameter transcription factor network. Mol Cell 51, 116-127.

Rual, J.-F., Ceron, J., Koreth, J., Hao, T., Nicot, A.-S., Hirozane-Kishikawa, T., Vandenhaute, J., Orkin, S.H., Hill, D.E., van den Heuvel, S., et al. (2004). Toward improving *Caenorhabditis elegans* phenome mapping with an ORFeome-based RNAi library. Genome Res 14, 2162-2168.

Shen-Orr, S.S., Milo, R., Mangan, S., and Alon, U. (2002). Network motifs in the transcriptional regulation network of *Escherichia coli*. Nat Genet 31, 64-68.

Watson, E., MacNeil, L.T., Arda, H.E., Zhu, L.J., and Walhout, A.J.M. (2013). Integration of metabolic and gene regulatory networks modulates the *C. elegans* dietary response. Cell 153, 253-266.

Watson, E., MacNeil, L.T., Ritter, A.D., Yilmaz, L.S., Rosebrock, A.P., Caudy, A.A., and Walhout, A.J.M. (2014). Interspecies systems biology uncovers metabolites affecting *C. elegans* gene expression and life history traits. Cell 156, 759-770.

Watson, E., Olin-Sandoval, V., Hoy, M.J., Li, C.-H., Louisse, T., Yao, V., Mori, A., Holdorf, A.D., Troyanskaya, O.G., Ralser, M., et al. (2016). Metabolic network rewiring of propionate flux compensates vitamin B12 deficiency in *C. elegans*. Elife 5, pii: e17670.

Watson, E., Yilmaz, L.S., and Walhout, A.J.M. (2015). Understanding metabolic regulation at a systems level: metabolite sensing, mathematical predictions and model organisms. Annu Rev Genet 49, 553-575.

Yilmaz, L.S., and Walhout, A.J. (2016). A *Caenorhabditis elegans g*enome-scale metabolic network model. Cell Syst 2, 297-311.

Yilmaz, L.S., and Walhout, A.J.M. (2014). Worms, bacteria and micronutrients: an elegant model of our diet. Trends Genet 30, 496-503.

Zhang, J., Holdorf, A.D., and Walhout, A.J. (2017). *C. elegans* and its bacterial diet as a model for systems-level understanding of host-microbiota interactions. Curr Opin Biotechnol 46, 74-80.

